# A weighted generative model of the human connectome

**DOI:** 10.1101/2023.06.23.546237

**Authors:** Danyal Akarca, Simona Schiavi, Jascha Achterberg, Sila Genc, Derek K. Jones, Duncan E. Astle

## Abstract

Probabilistic generative network models have offered an exciting window into the constraints governing the human connectome’s organization. In particular, they have highlighted the economic context of network formation and the special roles that physical geometry and self-similarity likely play in determining the connectome’s topology. However, a critical limitation of these models is that they do not consider the strength of anatomical connectivity between regions. This significantly limits their scope to answer neurobiological questions. The current work draws inspiration from the principle of redundancy reduction to develop a novel weighted generative network model. This weighted generative network model is a significant advance because it not only incorporates the theoretical advancements of previous models, but also has the ability to capture the dynamic strengthening or weakening of connections over time. Using a state-of-the-art Convex Optimization Modelling for Microstructure-Informed Tractography (COMMIT) approach, in a sample of children and adolescents (*n* = 88, aged 8 to 18 years), we show that this model can accurately approximate simultaneously the topology and edge-weights of the connectome (specifically, the MRI signal fraction attributed to axonal projections). We achieve this at both sparse and dense connectome densities. Generative model fits are comparable to, and in many cases better than, published findings simulating topology in the absence of weights. Our findings have implications for future research by providing new avenues for exploring normative developmental trends, models of neural computation and wider conceptual implications of the economics of connectomics supporting human functioning.

## Introduction

The study of the brain as a connectome using graph theory provides a powerful framework for understanding its computational and organizational principles^1,2^. There are well-characterized features of observable brain networks, such as its modular structure^3^, small-world propensity^4,5^, hierarchal organization^6,7^ and its geometric wiring structure^8^. Underlying these apparent features is the economic and energetic context in which brain network configurations exist^9,10^; preserving its physical, metabolic, and cellular resources while sustaining required neural function^9,11–14^. Due to intrinsic resource limitations for sustaining the brain’s organization, the connectome achieves a balance between the valuable connections required for appropriate functioning versus the costs of those connections to form, maintain and propagate signals^13–15^.

To better account for this complex organization, various flavors of probabilistic generative network model have been proposed since the early 2000s^12,16,17^. These models work by simulating the formation of connections in the brain in a step-wise fashion to produce synthetic connectomes. In essence, these models achieve compression in that they produce complex networks from just one or two tuned parameters^18^. Across studies, the systematic comparison of different parameter types, and tuning properties, highlights the fundamental constraints that govern the formation of a given network. When fit to empirical human brain data^17,19–21^, these models can shed light on the possible factors driving biological connectivity. For example, one emerging finding is that the preference for topological self-similarity, when modelled as a wiring rule (termed *homophily*), can approximate structural and functional connectome datasets across numerous species and scales (e.g.,^22,23^). This indicates that an important developmental principle may be that neural assemblies form connections with each other, based on how similar these assembles are to each other^24^.

In contrast with graph theoretical analyses of connectomes derived from *in vivo* diffusion magnetic resonance imaging (dMRI), which commonly consider the heterogeneous edge-weights that are observed^5,25,26^ (e.g., in terms of number of streamlines or fractional anisotropy), one major limitation of previous generative models is that they can only simulate the binary existence of connections in the connectome (i.e., reflected by a one or a zero corresponding to a connection existing or not, respectively). This means that they exclude consideration of connection weight heterogeneity^27^. This exclusion simplifies the engineering problem of simulating connectomes but significantly limits the scope of the scientific questions they can answer. First, because connectomics data have an intrinsic weighted structure, current generative network models largely ignore a potentially essential source of information. As a result, we may be missing insights critical for understanding the constraints that guide connectome formation. Second, the strength of relationships between regions (rather than just their existence) are crucial in neurocognitive development and highly sensitive to developmental change^28,29^. If generative models are to be useful for understanding this change they will need to capture weighted change. Third, in computational models that perform tasks (e.g., neural networks), weights mediate the extent to which errors propagate and facilitate computation. Without weights, it will be hard to integrate network formation and the computational capacities those networks afford (e.g., as in^30–33^).

We present a solution to these challenges through an extension of canonical generative models^16,17,20^ to a *weighted* generative network model of the human connectome. This model draws upon the economic insights from prior generative modelling^16,17^. However, we further extend the model, inspired by the principle of redundancy reduction^34^, but through the lens of network communication^35,36^, to account for how weights can adjust dynamically over time to optimize how signals are propagated across the brain’s connectome. Using state-of-the-art *in vivo* Convex Optimization Modelling for Microstructure-Informed Tractography (COMMIT)^37–39^ we demonstrate that this model is able to accurately approximate both the topology and weights of the human connectome. We provide potential future directions for the field and a framework for empirical findings may be incorporated into future models.

## Results

### The weighted generative network model

The weighted generative network model has two core algorithmic components driving the network’s developmental trajectory from its starting point to end state (**Fig. 1a**). The first is a binary generative network model^16,17^, in which connections *form* iteratively over time – a connection is generated where it previously did not exist. The second component is a weight optimization step, where connection strengths of existing connections *change* in a direction and magnitude to reduce communication redundancy in the connectome.

**Fig. 1.**
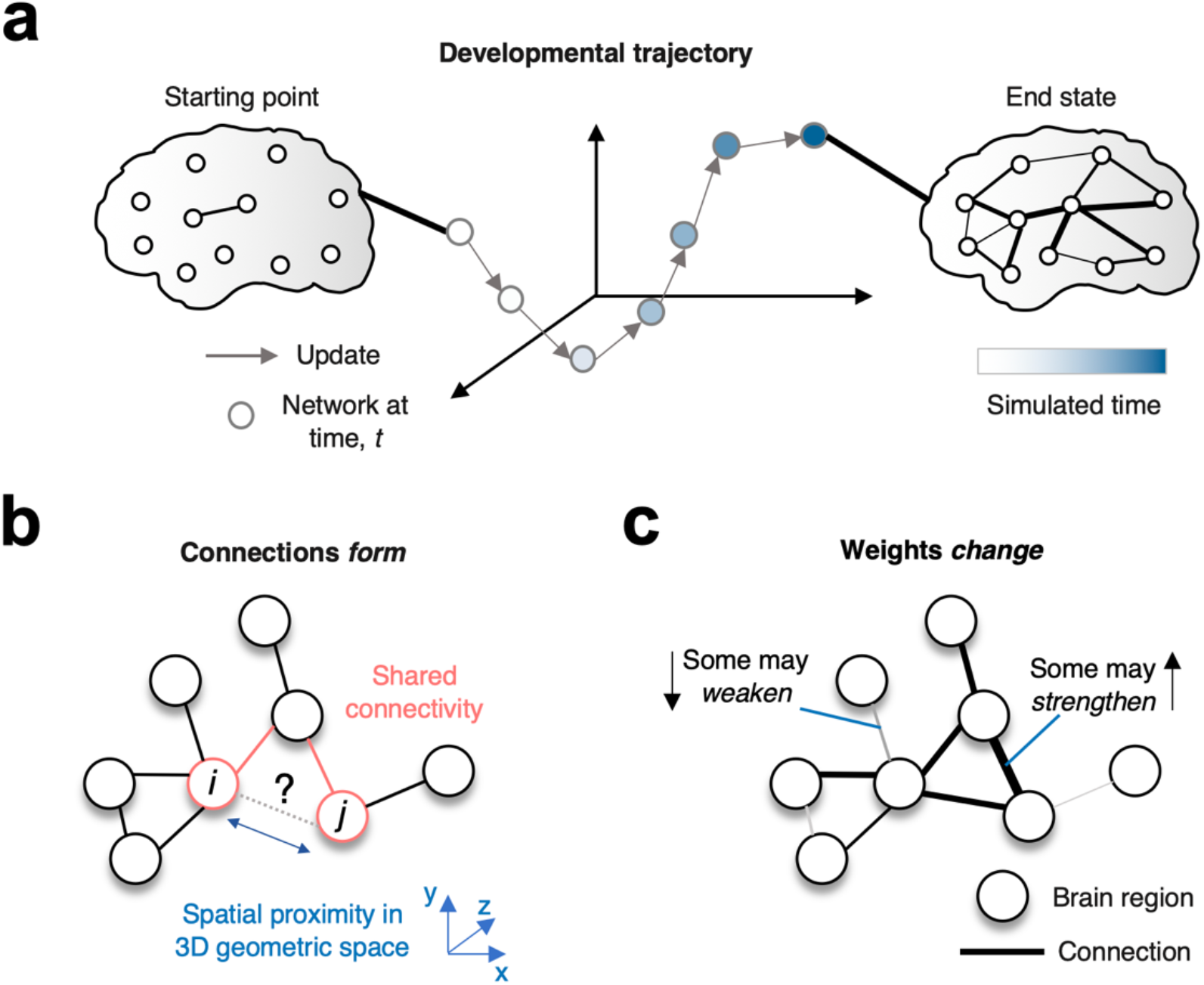
The network’s developmental trajectory comprises of connections forming and weights changing. **a** An illustration of the weighted generative process. As the simulated developmental time unfolds, the network moves through the feature space until it reaches its final destination. **b** In growing networks, connections form between regions. The information driving this process must be driven by factors outside the direct synaptic information present between the two regions, because this is absent. Two factors that could drive this are the current indirect connections linking the two regions or the spatial proximity of the regions. In this example, we highlight the shared connectivity (red) and spatial proximity (blue) between node *i* and *j*. An accurate model should demonstrate how connections form to generate connection topologies consistent with observations. **c** Connection weights change as some function of the presently available weights of the connections. An accurate model should demonstrate both weakening and strengthening over time of connections, that generates an organization of weights consistent with observations.

The distinction between connections *forming* (the first component) versus *changing* (the second component) in the model is not arbitrary. Before a connection is formed between two regions in the brain, each region does not have *direct* information from the other via a direct connection. Whatever information exists arises via other indirect connections (i.e., information passed via other, currently available, connections) or via some other non-synaptic means (e.g., paracrine signaling) (**Fig. 1b**). Once a connection has formed, we model changing connections as weights that change in a direction so as to reduce redundant communication. It may be that, as in developing neural systems, some weights strengthen and others weaken over time to achieve the goal of reducing unnecessary communication (**Fig. 1c**).

### Generative component 1 – Forming connections

Let’s consider the first algorithmic component: *forming connections*. For this we use the aforementioned generative network model. As stated previously, this model probabilistically adds a single connection according to the modelled costs and values afforded to the network^16,17^. The simulation stops when the number of connections mirrors the empirical network it is being compared to. It can be expressed as a simple wiring equation, updated over time:

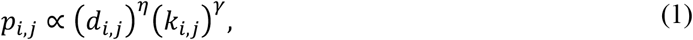

where *p*_*i,j*_ denotes the probability score of node *i* and *j* forming a connection. The algorithm has a winner-takes-all formalization such that a single connection is forced to form over the others, depending on this probability score at each iteration of the simulation over discrete time. *d*_*i,j*_ denotes the cost of wiring between node *i* and *j*. To prevent overfitting of the model introduced from pre-specifying fiber lengths (see *Discussion* for detail), we model this as the Euclidean distance between regions (node regions are defined in *Methods; MRI acquisition, processing and COMMIT*). In our sample (see *Methods; Participants*) the average correlation between fiber lengths and Euclidean distances for extant edges was *r* = 0.773 (SD 0.0119) corresponding to 59.7% explained variance. *η* is a parameter that determines the strength of the relationship between the cost of wiring and the probability of forming a connection. In empirical studies, best fitting models tend to show negative values, meaning that networks prefer shorter connections to longer connections, as measured by the Euclidean distance between two regions^16,17,20^. *k*_*i,j*_ denotes the topological value of forming a connection between node *i* and *j. γ* is the parameter that determines the strength of the relationship between the topological value and the probability of forming a connection.

The *k*_*i,j*_ term is given by an arbitrary topological relationship postulated *a priori* (also termed “wiring rule”). Prior work has shown that homophily (in particular, the *matching* rule) generative models robustly achieve the best model fits relative to other models^19–21^. Therefore, in order to make progress with the second component of the model, we focus only on this best performing homophily term for the first component, rather than cycling through all the various options. This matching rule computes the normalized shared connectivity profile – the average proportion of shared neighbours two regions have and has been used in numerous other studies to simulate the topology of empirical binary brain networks^12,19–21^. It is given by the following equation, where Γ_i_ where represents the set of node *i*’s neighbors:

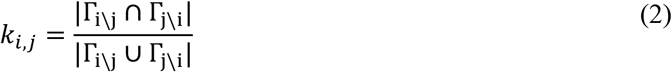

Where Γ_i\j_ is Γ_i_ but with *j* excluded from the set. ∩, from set theory, denotes the intersection of the neighbours (i.e., the overlap – in both sets). ∪, in contrast, denotes the union of the neighbours (i.e., the total set of neighbors from both sets). If there is a total overlap in neighbours, *k*_*i,j*_ = 1. If there is no overlap, *k*_*i,j*_ = 0. In summary, the formation of connections is modelled as a trade-off between the cost of forming a connection versus the topological value derived from having shared connectivity (under the matching rule).

### Generative component 2 – Changing weights

We now consider our second algorithmic component: *changing weights*. As the brain constructs itself, it does not simply add connections iteratively. Instead, as connections form, it simultaneously engages in continual plasticity, with some connection strengthened and others weakened over time^40^. But what drives this change over time? We propose a single optimization process that, as we later show, can simultaneously achieve the strengthening and weakening of connections: the weights of the network change to minimize its communication redundancy between its spatially-configured components. This idea stems from accounts of redundancy reduction as a core principle for economical sensory coding^34,41^ but through the lens of network communication^35,36^.

We will now outline how we operationalize redundancy in the context of communication. We define communication in terms of topological random diffusion of signals between regions on the weighted connectome^42,43^:

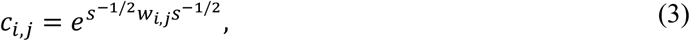

where *c*_*i,j*_ is the normalized weighted communicability between node *i* and *j*. This measure captures what proportion of signals that propagate randomly from node *i* would reach node *j* over an infinite time-horizon. It can be considered as equivalent to random diffusion or a random walk on the network graph. As such, it can be thought of as the extent to which node *i* and node *j* communicate. Here, *s* defines the diagonal matrix with the node strengths on the diagonal. *w*_*i,j*_ is the weighted matrix of the network, representing the strength of connections between nodes.

We use this operationalization of communication within an objective function, in which the growing network continuously updates its weights to minimize this evolving function. Crucially, in addition to topological paths constraining communication, distance is a key determinant of the timing of signal propagation in networks that may contribute to redundancy. Adding in these further distance considerations, we achieve the following objective function:

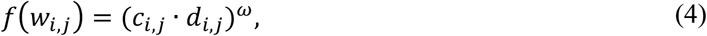

where *f(w*_*i,j*_) is the objective function that is calculated on the weight matrix *w*_*i,j*_. This takes in all learnable parameters (i.e., all non-zero elements of the weight matrix, *w*_*i,j*_). *d*_*i,j*_ is the Euclidean distance between node *i* and *j*, reflecting that the weights of longer connections are costly to maintain. *ω* is a parameter which varies the distribution of preference the network has to update weights. For example, when *ω* is a large positive value, it skews the optimization towards longer and more communicable edges. When *ω* is a small positive value, it softens this optimization disparity between edges. A similar term to this has also been used recently^44^.

Across the network, the goal is to minimize redundant communication in signals traversing physically in space. To achieve this optimization, at each time step *t* in the generative process we change the weights according to the following update rule:

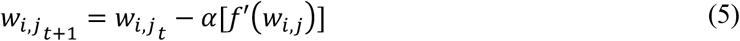

α is defined as the learning rate. The greater the learning rate the larger the jump in weight updates at each time point. *f*′(*w*_*i,j*_) is the first-order derivative of the objective function given in **Eqn. 4**, with respect to the network weights, *w*_*i,j*_. As previously stated, this has the effect of updating the weights of the network in a direction that minimizes communication redundancy in space. The first order derivative was estimated by simulating the objective function under small changes of individual weights (*δw*_*i,j*_)of 5% of the *w*_*i,j*_ value, taken incrementally five times, each in the positive and negative direction. The first order gradient is computed from these simulations, and weights are updated by the learning rate, *α*, at each timestep in the direction of the gradient. The sign of the update in **Eqn. 5** is negative because a positive gradient suggests that weights must be decreased to minimize redundancy (and vice versa, i.e., the subtraction facilitates the minimization of redundancy). For more detail as to the whole model algorithm, see Methods; *The weighted generative model algorithm*.

Once a weighted network was produced from the above process, we then assessed to what extent it mirrored empirical observations. We did this via an extensive model fitting procedure to compute model fit statistics called the *Energy*_*weighted*_ and *Energy*_*binary*_ which considered how well simulations approximated the empirical weights and topology respectively. Overall, the lower the energy value, the better the model fit. These energy statistics were calculated as the worst fit over a number of Kolmogorov-Smirnov (*KS*) statistics, which each measures the maximum distance between the cumulative density functions (CDFs) of some graph theory statistics in observed and simulated networks. To pick graph theory statistics, we extended those which have been used in prior work^12,19– 23^. For more detail, see *Methods; Model fitting*.

In **Fig. 2** we provide an illustration of the total weighted generative network modelling procedure to approximate empirical connectivity. To generate empirical connectomes, we used a *Convex Optimization Modelling for Microstructure-Informed Tractography* (COMMIT) approach^37,38^. COMMIT filters implausible streamlines from tractography and allowed us to assign the intra-axonal signal fraction (IASF) to each streamline (see *Methods; MRI acquisition, processing and COMMIT*). This provided us with a *in vivo* microstructural measure relating to measured axonal projection properties to provide a biologically meaningful measure of connection weights (**Fig. 2c**).

**Fig. 2.**
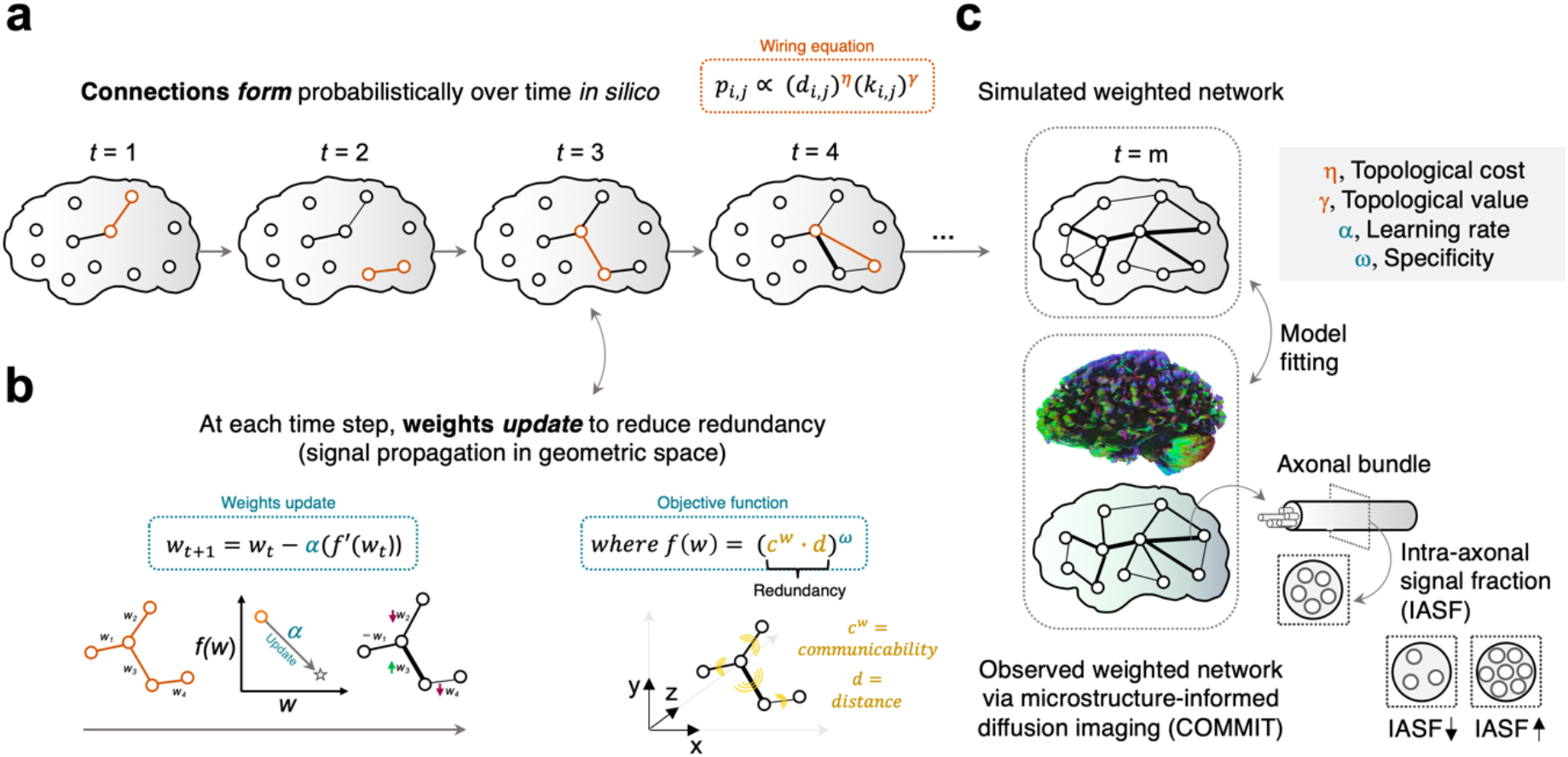
The weighted generative network modelling procedure. **a** Connections form probabilistically over time according to the canonical binary generative network model. **b** At each time point, once a connection has been formed, the network weights are optimized according to a learning rate, *α*, in a direction as to minimize the objective function *f(w*_*i,j*_) The first-order derivative *f*′(*w*_*i,j*_) is taken to do this. The objective function *f(w*_*i,j*_) here is defined in terms of the total network communicability, *c*_*i,j*_, and distance, *d*_*i,j*_. See *Methods; The weighted generative model algorithm* for detail of the whole generative process. **1. c** The simulation concludes when the number of connections is the same as the consensus empirical brain network. In the present work, we utilize microstructure-informed MRI which measures the intra-axonal signal fraction (IASF).

### Accurate simulation of weighted microstructure-informed connectomes

Through 3600 simulations of the above weighted generative network model, we charted the extent to which weighted connectomes could be simulated (**Fig. 3a**). In **Fig. 3b**, we show the *Energy*_*weighted*_ landscape as a function of the weight parameters, *α* and *ω*, at optimally fit *η* and *γ* parameters (see *Methods; Parameter selection*). As shown, the learning rate *α*and specificity term *ω* trade-off in the negative direction, such that low energy networks are generally found in the compromise between the two terms. **Supplementary Fig. 1** provides further landscapes, including the *Energy*_*binary*_

**Fig. 3.**
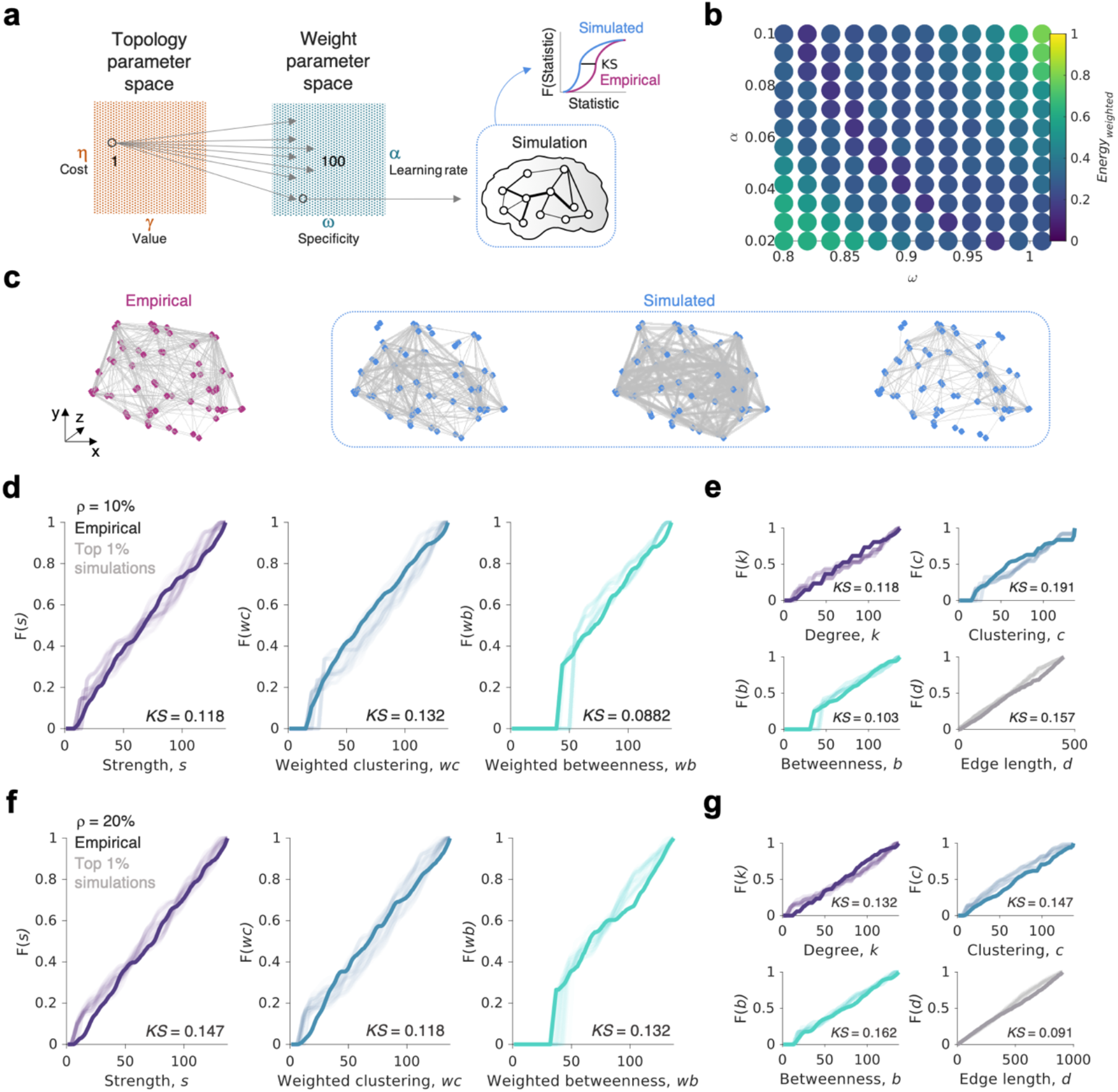
Simulating microstructure-informed connectome weights and topology simultaneously. **a** 3600 simulations were undertaken in total. This was done by sampling the topology parameter space from 36 locations spaced evenly apart (left) and running 100 simulations on each at regular intervals in the weight parameter space (middle). For each weighted network that was produced through this process, it was compared to empirical networks via the model fitting procedure (see *Methods; Model fitting*). This allowed for the determination of how well the model could approximate the empirical findings. **b** The *Energy*_*weighted*_ landscape (for sparse, ρ=10% networks) which depicts what combination of the learning rate, *α*, and specificity, *ω*, produce networks with low dissimilarity to observations. As described, each point entails 100 simulations with different combinations of *η, γ*. The best (i.e., minimum) *Energy*_*weighted*_ is given as the color (dark blue is low, yellow is high). **c** Observed (pink) and simulated (blue) weighted connectomes. The best (left), top 0.5% (middle) and top 1% (right) simulation is shown. **d** In sparse ρ = 10% networks, the cumulative distributions of strength (purple), weighted clustering coefficient (blue), weighted betweenness centrality (green). For each panel, the top 1% of simulations (36 in total) are shown in the lighter shade. The *KS* statistic is given only for the best performing simulation. **e** In sparse ρ = 10% networks, the cumulative distributions of degree (purple), clustering coefficient (blue), betweenness centrality (green) and edge length (grey). **f** The same as panel **d** but for denser ρ = 20% networks. **g** The same as panel **e** but for denser ρ = 20% networks.

We then sought to test our core question: to what extent is it possible to recapitulate both the topology and weights of empirical connectomes with a weighted generative network model? We first look at models fit to relatively sparse ρ = 10% networks. At this density, we found across our simulations, despite having to achieve more target features, the minimum energy concurrently achieved in weights and topology were comparable with the low values for binary networks: *Energy*_*weighted*_ of 0.157 (*KS*_*s*_ = 0.118, *KS*_*wc*_ = 0.132, *KS*_*wb*_ *=* 0.088, *KS*_*d*_ = 0.157) and *Energy*_*binary*_ of 0.191 (*KS*_*k*_ = 0.118, *KS*_*c*_ = 0.191, *KS*_*b*_ = 0.103, *KS*_*d*_ = 0.157). **Fig. 3c** and **Fig. 3d** show the cumulative density functions of simulated statistics compared to the empirical ρ = 10% network.

One criticism of our results is that, as with earlier work^19,20^, sparse networks may be easier to simulate accurately and achieve a good fit, simply because there are less connections to model. As such, we next aimed to replicate this finding in a denser ρ = 20% consensus network, containing twice the number of connections. We find highly similar results, with the weighted model actually doing a better job in most parts: *Energy*_*weighted*_ of 0.147 (*KS*_*s*_ = 0.147, *KS*_*wc*_ = 0.118, *KS*_*wb*_ *=* 0.132, *KS*_*d*_ = 0.091) and *Energy*_*binary*_ of 0.162 (*KS*_*k*_ = 0.132, *KS*_*c*_ = 0.147, *KS*_*b*_ = 0.162, *KS*_*d*_ = 0.091). **Fig. 3e** and **Fig. 3f** show the cumulative density functions of simulated statistics compared to the empirical ρ = 20% network.

How do the weighted simulations presented here compare to model fits attained from binary generative models? As one might expect, across our simulations, it is generally easier to simulate the network topology relative to being able to simulate the weights, with 96.1% and 98.8% of simulations showing a greater *Energy*_*weighted*_ relative to *Energy*_*binary*_ (respectively in sparse and dense consensus networks, **Supplementary Fig. 2a, b**). However, there are parallels in how the models fail to approximate the network statistics between weights and topology. In particular, prior findings have shown that binary homophily generative models struggle to approximate the clustering of the empirical observations and this is the part of the *Energy*_*binary*_ equation that tends to be worst approximated, reflected by being the highest *KS* statistic^19^. Here, we find a similar trend but for the weighted clustering measure, *KS*_*wc*_, which also generates the highest *KS* statistic in 83.1% and 92.3% of the simulations respectively in sparse and dense consensus networks (**Supplementary Fig. 2c, d**). **Supplementary Fig. 2e, f** show the broad relationship between the energy and *KS* statistics achieved through our modelling procedure.

### Evaluating models by their weighted and binary topological relationships

There is a large covariance between graph theory measures due to the dependencies between nodes via their connectivity^45^. For example, in empirical networks, it is common that regions with more connections tend to have connections with a higher average edge-weight^46^. Furthermore, due in part to the small-world propensity of brain network organization, regions which have high levels of clustered weights tend to have low betweenness centrality^5^. While some studies have examined this topological fingerprint more formally^22^, so far due to the lack of weighted information, this has been limited to the assessment of binary connections.

In **Fig. 4**, we show that while we have not explicitly simulated weighted generative networks to encompass these types of covariances, they arise as a result of the generative process. We find that simulations mirror (and slightly exaggerate) the dominant trend found in empirical networks (see **Supplementary Fig. 3** for the same findings on denser ρ = 20% networks). At first, these results may seem surprising because the weighted generative model explicitly detaches how connections form from how weights change (see *Methods; The weighted generative algorithm*). However, as we will outline in the next section, while there is a distinction between how weights and topology occur algorithmically, the principle of redundancy reduction means that topology constrains how weights arrange in a direction that aligns with empirical data. Put simply, despite the computational separation of connection formation from weight change, one will shape the other.

**Fig. 4.**
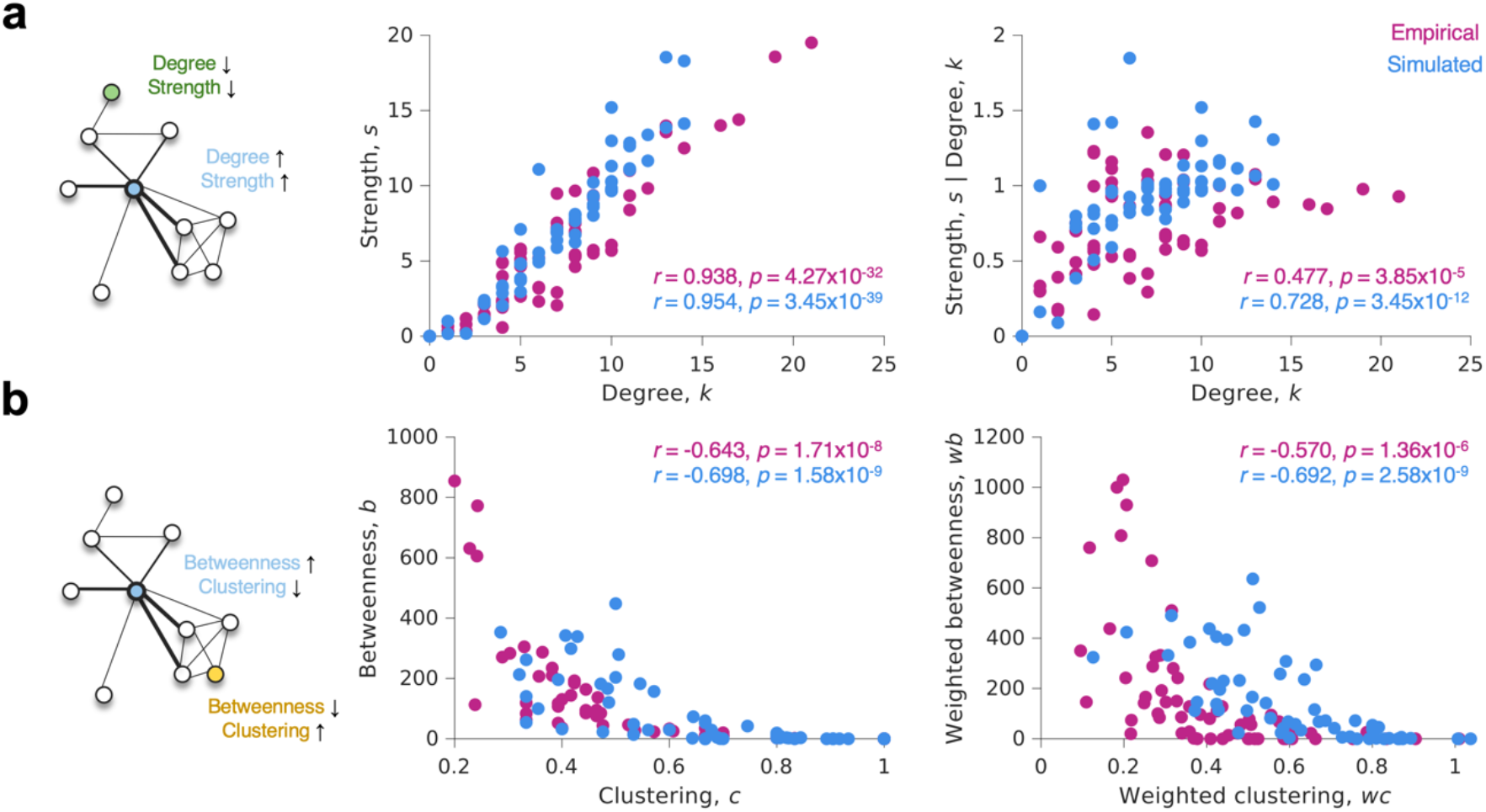
Topological relationships between weighted and binary network statistics in empirical and simulated connectomes. **a** In canonical idealized brain networks (left), regions with high numbers of connections also have stronger connections (light blue node) and vice-versa (green node). We show that in the best simulated networks, this is also the case in terms of the relationship between the number of connections of region has and the strength of those connections (middle). To ensure that we control for the analytical relationship between degree and strength, we also provide a version here the degree is controlled for in the strength measure (right). **b** In canonical idealized brain networks (left), regions with high levels of clustering have lower levels of betweenness centrality (yellow node) and vice-versa (light blue node). We show that in the best simulated networks, this is also the case in terms of the relationship clustering and betweenness (middle). We show this also for the weighted versions of the measure (right). All findings are given for ρ = 10% networks (see **Supplementary Fig. 3** for ρ = 20% networks).

### Communication redundancy reduction and the weighted connectome

So far, we have shown that by developing *in silico* networks which form incrementally according to self-similarity (i.e., homophily) in space, while concurrently optimizing weights to minimize communication redundancy in space, it is possible to produce macroscopic networks with high statistical similarity to observations. Although we have outlined the logic underpinning our formulation of the weighted generative algorithm, we have not directly provided an account for precisely *why* the model successfully fits the data.

Redundancy reduction is a core neuroscience principle dating to the 1960s^34^ where Barlow hypothesized that the goal of sensory processing was to recode redundant sensory inputs into a factorial code with statistically independent components^41^. This idea has since inspired numerous learning algorithms^47–49^. The current work, rather than focusing on redundancy in sensory processing, focusses on redundancy in terms of how regions themselves propagate signals between each other.

For example, a network that has a lot of communication is likely to be redundantly communicating an abundance of signals between numerous regions. In contrast, a network with little communication is required to communicate efficiently its relatively sparse signals between regions. This way of considering communication redundancy is consistent with an efficient coding framework, which proposes that the brain transmits maximal information in a metabolically economical or compressed form to improve future behavior^36,50^. By operationalizing this mathematically in **Eqn. 3** (as in analogous work^51^) we have defined a type of redundancy that is minimized throughout the generative process.

How does this principle of redundancy reduction in communication lead to our empirical observations of connectome organization? To examine this question, we conducted the following experiment. We undertook the same optimization process in the weighted generative model, but carefully evaluated how redundancy changes as a function of individual weights changing over time. We depict the main findings of this experiment in **Fig. 5a**. Starting from a simple exemplar binary network of nine nodes, we compute how changing individual weights in small increments of 2.5% in the positive and negative direction (*δw*_*i,j*_) changes the total level of communication (*ΣC*) in the network (as computed from **Eqn. 3**). As shown, not all changes of weights cause the same effect: as some connections are strengthened communication decreases, but in others, you must weaken connections to get the same trend of communication decrease.

**Fig. 5.**
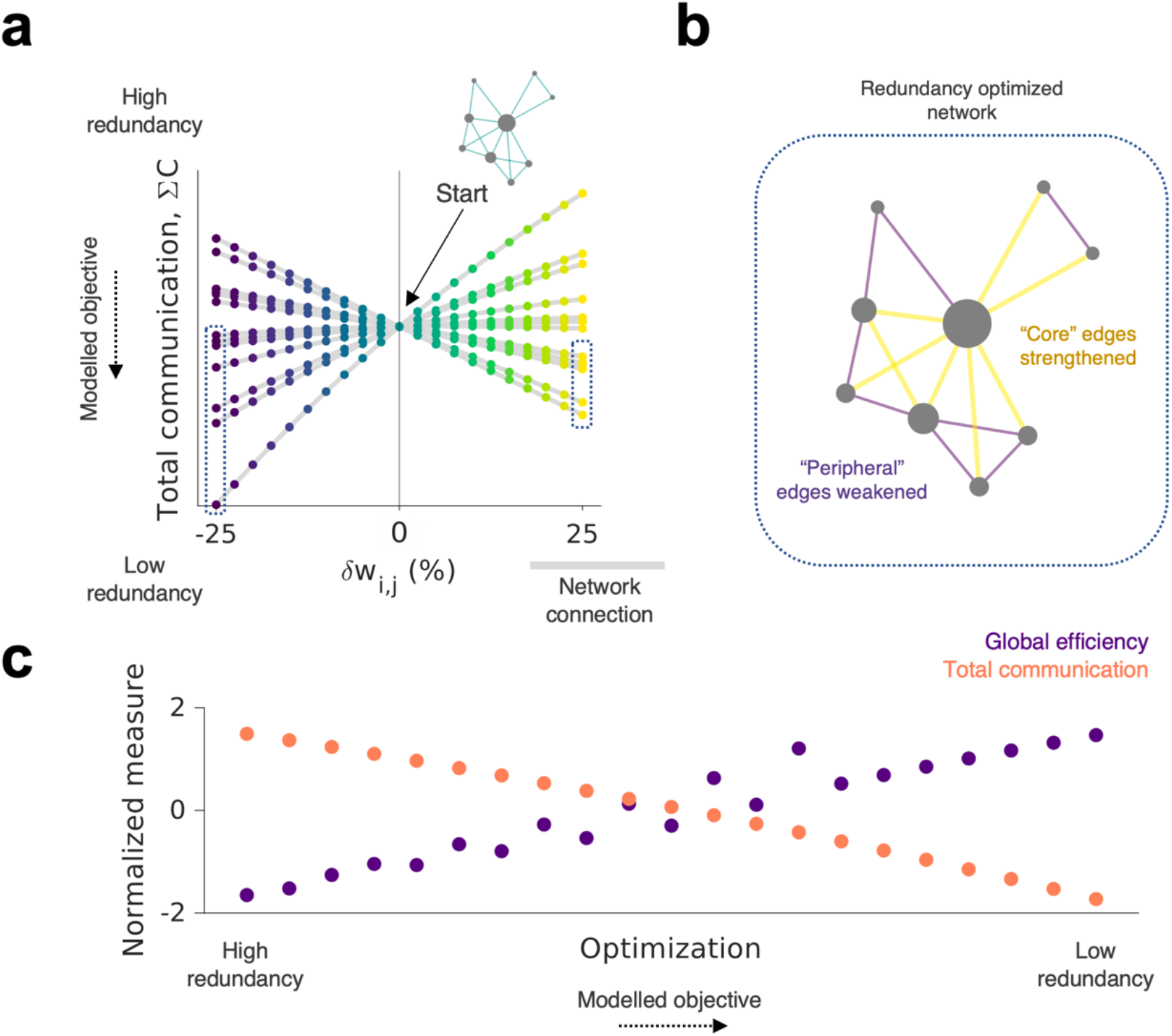
Redundancy reduction leads to an efficient patterning of connectivity weights. **a** Starting from the given binary network, we incrementally strengthened or weakened the edge weights (*δw*_*i,j*_) in increments of 2.5%. At each change, we recorded the total communication across the network (*ΣC*) which is a measure of redundant communication existing in the network. Each line represents a different edge in the network (a total of 16 edges) as the connection is changed. As shown, there is a “crossing” phenomenon, where strengthening or weakening of connections cause opposite effects on the redundancy present in the network. As highlighted, some connections achieve minimal redundancy when the connection has been weakened (left, blue dashed box) but others achieve this when they have been strengthened (right, blue dashed box). **b** There is a topological relationship between where the connection is in the network and its relationship between connection weights and redundancy, such that to achieve a redundancy optimized network you must strengthen core connections but weaken peripheral connections. **c** This phenomenon has the effect of causing the weights to become strengthened in the core of the network – equivalent to greater integration – causing greater efficiency (purple) but reduced communication (orange).

This “crossing” phenomenon (as seen in **Fig. 5a**) can be explained by the fact that communication redundancy is minimized when core connections – that are topologically central to information flow – are strengthened, but peripheral edges are weakened (visualized in **Fig. 5b**). In summary, the process of redundancy reduction leads to the bottlenecking of signal propagations within relatively few core connections, consequently leading to an increased efficiency in regional communication (**Fig. 5c**). This allows for the network to prioritize the flow of communication through topologically central nodes, allowing for efficient integration of communication across the connectome.

## Discussion

### Redundancy reduction in network communication

One key finding is that by reducing communication redundancy, it is possible to approximate both connectome topology and weights. There are many other ways, in principle, that the goal of this algorithm could have been instantiated. For example, one could imagine an alternative multi-step algorithm in which connections are added and/or then removed in sequence at each time-step. However, the present approach provides several major benefits relative to such a solution. First, in the current model the strengthening and weakening of connections can be accounted for via a single optimization process depending only on the communication redundancy within the network. Achieving these heterogeneous magnitudes and directions of weight changes over time (i.e., both strengthening and weakening) is not trivial, particularly when there is no supervision in the learning process. Second, our optimization process is theory-driven, rather than mathematically arbitrary. The idea that redundancy should be constrained in biological systems is highly congruent with multiple theoretical perspectives in neuroscience^34,41,52,53^. In summary, not only does this model solve a somewhat challenging engineering problem, but it does so in a way that resonates with biological theory.

The way we formalize redundancy reduction here is not identical to how the original efficient coding hypothesis framed redundancy. The original hypothesis concerned how the goal of sensory processing was to recode redundant sensory inputs into a code with statistically independent components, as to remove redundant signals from external stimuli within the internal representations^34^. Here, rather than focusing on the redundancy within internal representations, we focus on redundancy within the *communication* of the network. This work draws parallels between information theory accounts of neural communication via compression in the connectome^36^ but through the lens of resource rationality^54,55^: where each node in the graph, as it develops, aims expend the least possible amount of communication expenditure^13,56^. Moreover, we demonstrate a particularly interesting observation that the reduction of redundant communication can account for how, over time, networks may incrementally integrate their weighted connections – an observation entirely congruent with studies examining topological changes throughout child development^57^.

### In context of prior structural generative network model findings

To date, generative models have highlighted numerous insights regarding connectome organization ^18,58^. In particular, cost-minimizing homophily models (as used within the current work) have been shown to quite consistently generate realistic connectome topologies across a range of scales, species and modalities^12,17,19,21,23^ (although see^59^). Wiring parameters have been shown to link to cognition^19,21^, age^19,20^, polygenic risk for Schizophrenia^21^ and adversity in a rodent model^23^. As our weighted models builds on this foundation, we expect that theoretical questions they can answer may extend and compliment this previous work.

One way it may do this by capturing more biologically meaningful parameters which relate to the organization of connectome edge-weights. For example, the parameter controlling wiring length, η, has been shown to correlate with polygenic risk for Schizophrenia^21^ – where subjects with higher scores tended to have a lower magnitude negative η suggesting a softer cost penalty on connectivity. A weighted model may be able to elucidate more specific weight-topology interactions that may play a role in disease onset or altered development^58^. It may be able to better elucidate age-related changes shown in lifespan data^20^ or how weighted connectivity early in development^60^ may be modelled in combination with genomic^12^ or cytoarchitectural^61^ data.

Another way weighted models may extend our analysis is by informing how topology and weights both interact to provide computationally efficient networks able to perform computation^30–32,44^. For example, recurrent (task-solving) neural networks have been shown to develop brain-like topological features through a very similar optimization procedure described in the current work^44^. In this network in particular, we highlight the interplay between a growing topological network (via a homophily rule) which subsequently shapes how weights organize themselves through bottlenecking of weights within topological core regions of the network (see **Fig. 5**). We anticipate this observation will lead to a number of new theoretical insights at the intersection of network neuroscience and neural network research^24,62^.

### Limitations and future research

Below we list numerous limitations of the present study and point to how these can be reasonably mitigated in future research:

#### Computational expense

On a typical desktop computer, binary generative models take approximately one second to compute a binary generative model (ρ= 10% connectome, 227 connections). However, our weighted models take approximately 300 seconds (∼300x slower). This is for two main reasons. The first is that computing the first-order derivative of our objective function (**Eqn. 4**) becomes increasingly difficult as the network grows. This leads to an intrinsic slowing down of the model over the network’s formation. The second more important factor is that in the present study we compute the gradient manually through a model-based simulation of the objective function with respect to the weights. This is computationally expensive, but future work will be able to mitigate this through employing faster approaches to compute derivatives.

#### Consensus model fits rather than individual subjects

As there is a large computational expense for computing the current models, it limited our ability to accurately fit models to individual subjects. This leaves numerous open questions: how do weight parameters vary across individuals? Do these models better map to measures of cognitive performance or polygenic risk? While we suspect they will, we cannot yet confirm our findings will generalize to individual subjects robustly. It also may be that our parameter search would need to be widened to encompass individual subjects. An extended approach similar to the fast landscape generation method^63^ would be particularly helpful to approximate individual subject accurate parameter estimates.

#### Parcellation coarseness

We limited our analysis to the 68-node DK parcellation which, although studied before with generative models^19^, is a coarse parcellation. It is unknown how these results will generalize to finer resolutions such as the Brainnetome^64^ or Schaefer^65^ atlases. Another effect of the parcellation may reside in the inter-node distance distributions. For example, more heterogeneously spaced parcellations will likely more easily generate modular networks simply by virtue of the *a priori* locations of the regions. Our current study is limited by not exploring these effects, but future work can explore this.

#### Negative weights

Numerous studies have used functional, rather than structural connectomes, when using binary generative models^17,22^. While communication models have been argued to allow for better mappings between structural and functional modalities^35^, our model does not deal with negative weights, which is intrinsic to correlations. This leads to a natural fit between our model and structural data, which naturally contains non-negative edge-weights. Of note, other generative models which can capture weights, in the form of stochastic blockmodels (which are useful characteristic network community structure), can deal with negative edge weights^66^.

#### Wiring rules

In context of prior work, we only looked into the homophily model. However, given that topology is an influence on weighed optimization process, we think it is possible that other rules will yield subtly different results. Future work should look to explore how weights differentially configure themselves in context of different connection formation rules.

#### Other constraints

As in most other studies^20,21^, we use Euclidean distance as a measure of cost of connection formation guiding topology and weight change. In contrast with fiber length constraints, this has the benefit of removing any *a priori* limits to how the network’s topology can be generated. This is because fiber length data only exists for extant connections but Euclidean distances can be computed between all nodes. However, in this study approximately 60% of variance in fiber lengths of extant connections can be explained by the Euclidean distance – representing a relatively large variance explained (see^20^ for comparisons). Adding fiber lengths as a constraint will, on one hand, reduce the search-space for simulations that mirror observations but, on the other, may reveal more specific generative principles specific to our observations. Moreover, while distance is a key determinant of signal propagation (and hence influences weight change in our model), so too are other factors we hitherto do not model such as axon diameter or the g-ratio^67^. Other constraints could be added into the model, such as cytoarchitectural, gene-expression or receptor-expression similarity^68–70^.

#### Seeding and connection formation weighting

This study initializes the formation of the network from an empty network so that no prior information is given to the network. In practice, this means that development starts where the connectivity cost is least, which is by definition in the center of the space. This early initialization will have a key effect on the model^71^ but is not very biologically plausible (see^72,73^). Aside from the earliest simple generative models^16^, this fact been largely ignored. Future work should address this by systematically testing how initial network conditions influence the resulting simulation.

## Conclusions

We present a new weighted generative network model, capable of capturing the weighted topology of the human connectome. This model solves a major limitation of prior research, principally because it is weighted, extending our capability to accurately approximate both the weights and topology of the connectome. We introduce several novel contributions, including an openly-available function that can be used to simulate empirical neuroscience data, a demonstration of this model applied to microstructure-informed tractography data (COMMIT), in addition to a principled mechanism for explaining why weights become configured as they do via a principle of communication redundancy reduction. By using this model, we extend the economic accounts of brain organization, providing a better understanding of how the brain may not only balance the valuable connections necessary for appropriate functioning with metabolic costs, but also how their weights may be modified, in context of the topology, to minimize redundant communication as it forms.

## Acknowledgements

Danyal Akarca, Jascha Achterberg and Duncan Astle are supported by UKRI MRC funding (MC-A0606-5PQ41). Danyal Akarca and Duncan Astle are funded by the James S. McDonnell Foundation Opportunity Award for Understanding Human Cognition and the Templeton World Charity Foundation, Inc. (funder DOI 501100011730) under the grant TWCF-2022-30510. Jascha Achterberg receives a Gates Cambridge Scholarship. Jascha Achterberg was an intern at Intel Labs at the time of writing. The data were acquired at the UK *National Facility for In Vivo MR Imaging of Human Tissue Microstructure* funded by the EPSRC (grant EP/M029778/1), and The Wolfson Foundation. This research was funded in whole, or in part, by a Wellcome Trust Investigator Award (096646/Z/11/Z) and a Wellcome Trust Strategic Award (104943/Z/14/Z). For the purpose of open access, the author has applied a CC BY public copyright licence to any Author Accepted Manuscript version arising from this submission.

## Methods

### Participants

Our main cohort contains *n* = 88 total participants (mean age = 12.56 years, SD age = 2.94 years, minimum age = 8.01 years, maximum age=18.96 years). The sample contains *n* = 46 girls (mean age = 13.23 years, SD age = 3.13 years, minimum age = 8.01 years, maximum age = 18.96 years) and *n* = 42 boys (mean age = 11.82 years, SD age = 2.57 years, minimum age = 8.03 years, maximum age = 16.77 years). There is a slight interaction between age and sex, whereby the girls in our cohort were older (*p* = 0.025) (**Supplementary Fig. 4**).

### MRI acquisition, processing and COMMIT

All *n* = 88 participants were scanned on a 3T Siemens Connectom system with ultra-strong (300mT/m) gradients, using: multi-shell diffusion magnetic resonance imaging (dMRI) acquisition (*TE*/*TR* = 59/3000 ms; resolution 2×2×2 mm^3^) with *b* ∈{500, 1200, 2400, 4000, 6000} s/mm^2^ in 30,30,60,60,60 directions, respectively and additional 14 *b* = 0s/mm^2^ interleaved volumes; 3D MPRAGE (*TE*/*TR* = 2/2300ms; resolution 1×1×1mm^3^). dMRI data were pre-processed as outlined elsewhere^74^.

To provide a more ‘biologically-informative’ assessment of brain connectivity, we used a *Convex Optimization Modelling for Microstructure-Informed Tractography* (COMMIT) approach^37,38^. Briefly, COMMIT deconvolves specific microstructural features on each fiber to recover individual streamline contributions to the measured signal. To achieve this, we performed multi-shell multitissue constrained spherical deconvolution (MSMT-CSD) and generated a whole-brain probabilistic tractogram seeding from the white matter to generate 3 million streamlines. We then applied COMMIT with a stick-zeppelin-ball model^75^ to simultaneously filter out implausible streamlines and assign an intra-axonal signal fraction (IASF) to each one. Thus, for all subjects we set the following diffusivities *d*_*par*_ = *d*_*parzep*_ = 1.7 × 10^−3^ mm^2^/s, *d*_*perp*_ = 0.6×10^−3^ mm^2^/s, *diso* ∈{1.7, 3.0}×10^−3^ mm^2^/s^76^.

Connectomes were subsequently built using the FreeSurfer Desikan-Killiany (DK) parcellation as nodes (68 cortical) and by then assigning the total IASF associated to each bundle as edge-weights^39^.

### Group consensus thresholding

As the modeling approach given is highly computationally expensive (taking ∼300x more computational time than the binary model alone), rather than fitting our models to each of the *n* = 88 participants we performed our modelling procedure on a consensus network built from the *n* = 88 sample. Utilizing a consensus also reduced the impact of false positives, false negatives^77^ and any effect of inconsistencies in the reconstruction of subject-level connectomes^78^. We generated the group-level consensus networks from the sample level IASF-weighted connectomes, which had a thresholded mean density of ρ = 34.5%. We provided absolute thresholds of 0.1839 and 0.0467 to these IASF-weighted networks to enforce a density of both ρ = 10% and ρ = 20% across the sample, before running the consensus procedure. These densities were picked to best replicate the literature, which has commonly used ρ = 10% or ρ = 20% networks^12,20,22^ but more importantly so that we can establish any effects of the models on relative sparse versus dense networks.

To generate an accurate group-level representative consensus, we used *fcn_group_bins()* in Matlab 2020b, which has been comprehensively detailed elsewhere constraining node-to-node distances by the node-centroid Euclidean distances^79^. This approach retains the topological characteristics of individual subject networks and preserve within-/between-hemisphere connection length distributions of the individual participants.

The result of this procedure were two binary graphs (ρ= 10% and ρ = 20%), which acted as the observed group topological consensus network. We then used the mean IASF weights across all participants as the attributed weighted edges to complete the consensus weighed network. These consensus networks contained 227 and 454 connections respectively across the 68-node DK parcellation. See **Supplementary Fig. 5** for more detail of these consensus networks, including their network statistics. All network statistics were computed using the Brain Connectivity Toolbox (BCT)^80^.

### The weighted generative model algorithm

In this work, we construct simulated networks using a weighted generative network model, extending prior work^16,17,19,20^, to additionally encompass weights. We described the approach in earlier sections (see *Results; The weighted generative network model*) but we additionally provide a step-by-step algorithm here.

The algorithm begins from some initial starting condition. Here, we initialize the network as empty (i.e., zero connections) within the 68 cortical node DK parcellation scheme.

Edge connections are added in a highly analogous way to previous work which employs the canonical generative network model (see^20^ for further detail). Connections are added one at a time (i.e., connections *form*) over a series of steps until *m* total connections are placed. As stated in the above section, the *m* was computed as a group consensus over different controlled densities, leading to *m* = 227 and *m* = 454 (ρ= 10% and ρ = 20% respectively). This meant that the simulation achieved the same number of connections as the empirical data.

At each step, we allow for the possibility that any pair of presently unconnected nodes, *i* and *j*, to become connected. But this happens probabilistically, such that the relative probability score is given by **Eqn. 1**. As described in **Eqn. 1**, this is governed by a trade-off between the wiring cost (determined via Euclidean distances) and the homophily matching rule, which demarcates the topological overlap in connectivity of two nodes (given in **Eqn. 2**). We provide some extended reasoning for this part of the algorithm (see *Results; Generative component 1 – forming connections*).

At each point in which a connection is added (e.g., see **Fig. 2**) we take some property of the network and change the weights of the network, incrementally, in a direction as to partially minimize this property. This property (also termed objective function, *f(w*_*i,j*_)) is defined in **Eqn. 4** and is computed as the combined total weighted communicability (**Eqn. 3**) multiplied by the Euclidean distances present in the network. We provide our reasoning for this in terms of communicative redundancy reduction (see *Results; Generative component 2 – changing weights*). The *ω*term changes the specificity of the objective function to specific weights, such that the closer *ω*tends to zero the more equally distributed the weight changes are across the network over its simulated development. The greater *ω*becomes in the positive direction, the greater it emphasizes changes to weights that contribute to highly communicable and physically distinct connections.

To change the weights at each time step, we compute the first order derivative of the objective function, *f*′(*w*_*i,j*_,) which calculates an estimated gradient for each edge-weight must move to achieve the objective. Some weights are strengthened and some are weakened in this process. We then update the weights in this direction according to the update rule given in **Eqn. 5**, by some magnitude, otherwise termed learning rate, *α*. The greater *α*is, the greater that weights change at each time point after a new edge is added.

### Model fitting

In binary generative modelling work, numerous model-fitting functions have been proposed that assess network statistics^20^ or their statistical correlations^22,81^. To ensure we are fitting both the topology and the weights of the network, we simultaneously assessed (i) the binary representation of the produced network using a documented binary energy equation^20^ and (ii) a weighted version of the same energy equation, where weighted versions of the same graph measures are considered. These are given respectively in the following equations:

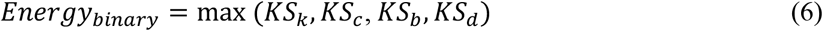

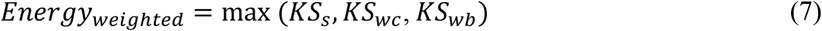

*KS* is the Kolmogorov-Smirnov statistic, defined as maximum difference between the empirical and simulated cumulative density function of the graph theory statistic. As a result, both the *Energy*_*binary*_and *Energy*_*weighted*_ can be thought of taking the worst of the measured comparisons. If the model fit remains low, the fit must necessarily be the same or lower across all considered statistics. The *Energy*_*binary*_ equation considers the node degree, *k*, clustering coefficient, *c*, betweenness centrality *b*, and edge lengths, *d*. The *Energy*_*weighted*_ equation considers the node strength, *s*, weighted clustering coefficient, *wc*, betweenness centrality *wb*. The edge length is not considered again because it was captured in the **Eqn. 6**. In all cases, we report the simultaneous model fits for both *Energy*_*binary*_ and *Energy*_*weighted*_ To enable comparability of intra-axonal signal fraction (IASF)-weighted connectomes, we normalized all connectomes using BCT’s *weight_conversion()* function^80^.

### Parameter selection

Our weighted generative algorithm contains four free parameters: *η, γ, α* and *ω*. The first two relate to the formation of connections within a binary model: *η* (connection length) and *γ* (topological value), and have been previously documented under a matching homophily rule to approximate networks accurately in the range of moderately negative *η* scalar values and positive *γ* scalar values slightly above zero^19,20^. Following some trial tests, we established an approximate window of -3.7 < *η* < -2.7 and 0.35 < *γ* < 0.40 for which we undertook more thorough parameter fitting. As the generative algorithm detaches the connection formation *η, γ* parameters from *α, ω* weight update parameters, we used these previously reported ranges of *η* and *γ* to significantly reduce our computational burden. We subsequently conducted a parameter grid-search across *α* (the weight learning rate) and *ω* (connection optimization specificity) to examine to what extent the weighted generative model could approximate both the topology and weights within these parameter windows. Following basic exploration, we conducted our search in the range of 0.02 < *α* < 0.1 and 0.85 < *ω* < 1.05. We fit our models to consensus IASF brain networks, derived at a density of both ρ = 10% and ρ = 20% to observe effects of numbers of connections on the network (see *Methods; COMMIT Signal fraction & Methods; Group network consensus procedure*). A total of 3600 simulations were run on these networks (total 7200) to fit the four parameters. All analyses were conducted with no seed network.

## Data availability

Derived MRI outputs can be made available upon request. Generative model outputs will become available on the Open Science Framework upon publication. A pointer to these will become available at https://github.com/DanAkarca/weighted_generative_models.

## Code availability

The weighted generative model function is available for open use at: https://github.com/DanAkarca/weighted_generative_models. All code to replicate the present study will become available at the same repository, upon publication. The code we used run COMMIT is available at https://github.com/daducci/COMMIT.

