## Supplementary Materials for "A weighted generative model of the human connectome"

### Contents

**Supplementary Fig. 1.**
Energy model landscapes for binary and weight model fits across all tested parameter combinations.

**Supplementary Fig. 2.**
The relative model fits for simulating connectome topology versus edge-weights.

**Supplementary Fig. 3.**
Replication of simulated topological relationships, in dense networks.

**Supplementary Fig. 4.**
Sample age and sex characteristics.

**Supplementary Fig. 5.**
Empirical consensus network statistics.

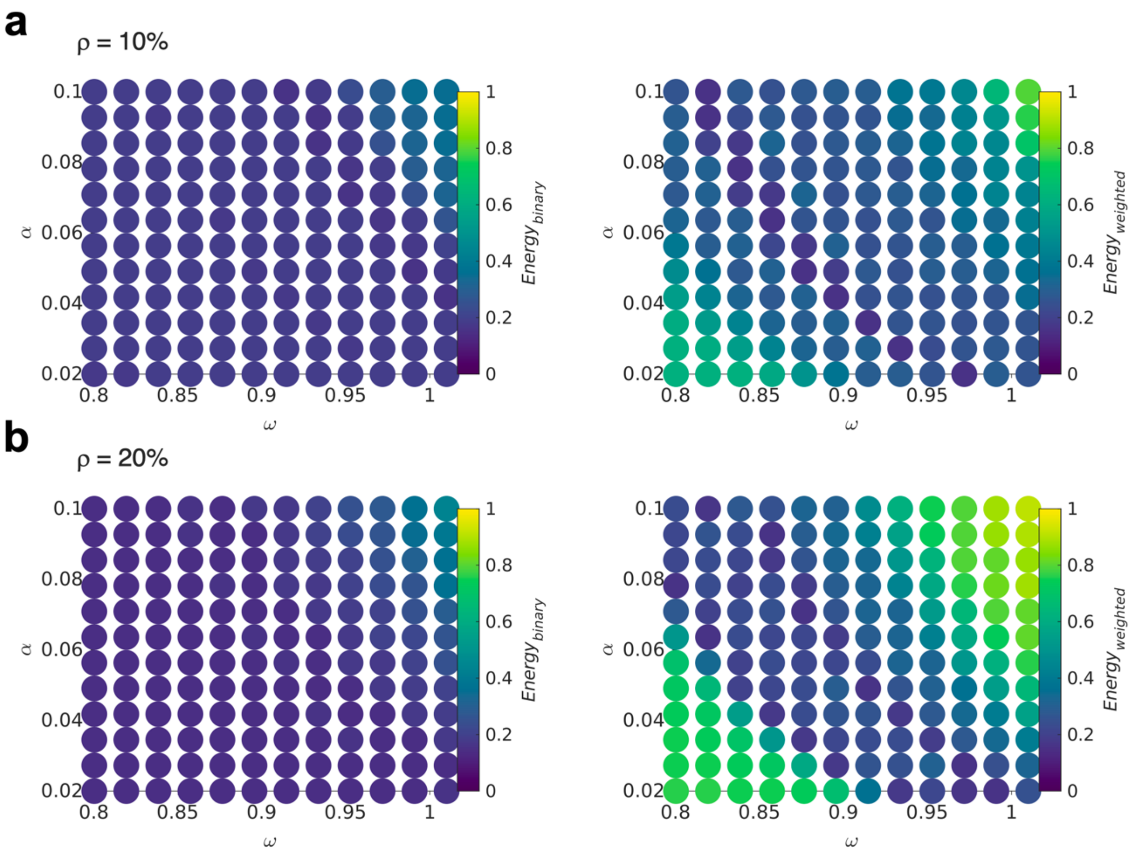


**Supplementary Fig. 1 | Energy model landscapes for binary and weight model fits across all tested parameter combinations. a** The $Energy_{binary}$ (left) and $Energy_{weighted}$ (right) landscapes for sparse (ρ = 10% density networks) which depicts what combination of the learning rate, $\alpha$, and specificity, $\omega$, produce networks with low dissimilarity to observations. The right panel is the same as depicted in **Fig. 3b**. **b** The same but for dense networks (ρ = 10% density networks).

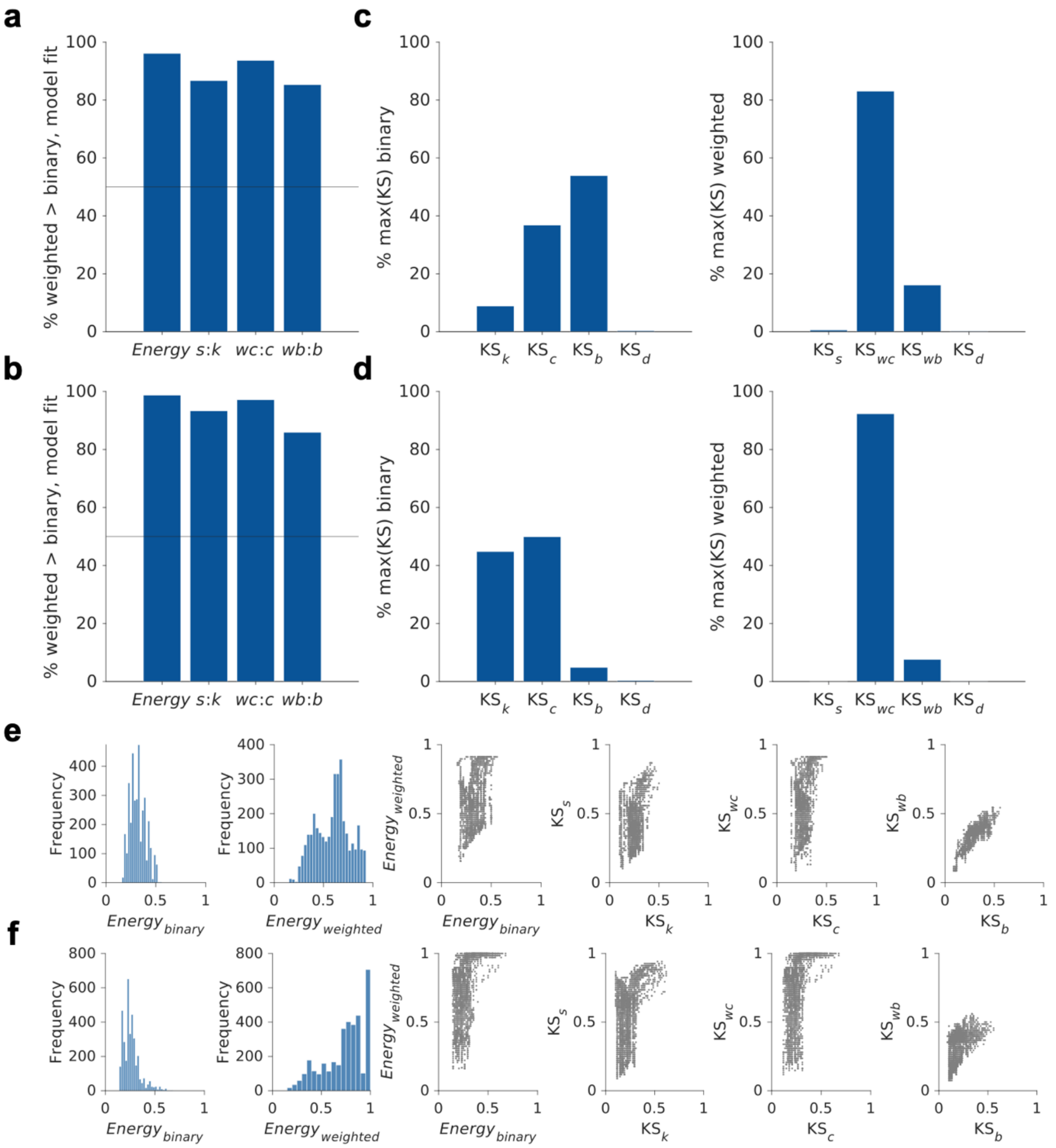


**Supplementary Fig. 2 | The relative model fits for simulating connectome topology versus edge weights.** **a** For sparse (ρ = 10% density) networks, we show the percentage of simulations in which the weighted model fit statistics was greater than the binary model fit statistic. A line is drawn at 50% which would suggest that they do equally well. In all comparisons, the weighted model fit is greater in magnitude that the binary, suggesting a worse performance all round. **b** The same is shown but for dense (ρ = 20% density) networks. **c** For sparse networks, we show what *KS* statistic determined the energy value (which is computed as the maximum, i.e., worse, of the four comparisons). We show this for binary ($Energy_{binary}$, left) and weighted ($Energy_{weighted}$, right). As shown, in sparse networks it is the betweenness centrality that mostly limits binary model fits (left) but the weighted clustering which limits weighted model fits (right). **d** The same plots are shown for dense networks. In contrast to sparse networks, it is clustering and the node degree which limits binary model fits (left). However, the same trend is found in terms of weighted clustering limiting weighted model fits (right). **e** Histograms and scatter plots describing the distribution of model fits and the relationships between binary and weighted *KS* statistics in sparse networks and **f** dense networks.

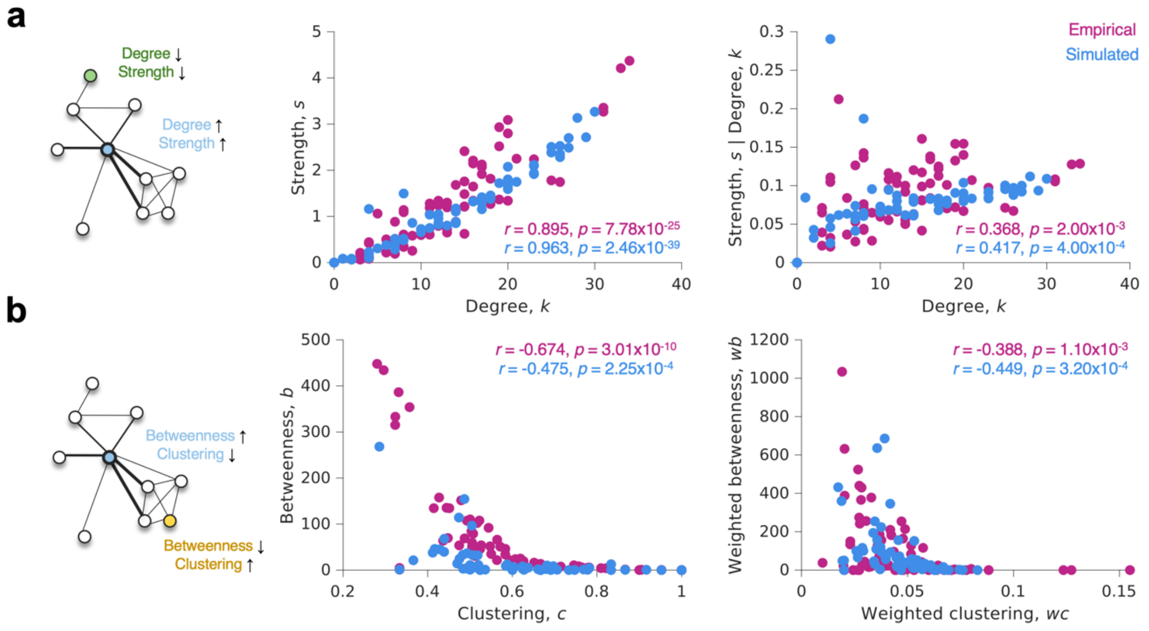


**Supplementary Fig. 3 | Replication of simulated topological relationships, in dense networks.** This is the same as **Fig. 4** but shown for dense (ρ = 20% density) networks. **a** In canonical idealized brain networks (left), regions with high numbers of connections also have stronger connections (light blue node) and vice-versa (green node). We show that in the best simulated networks, this is also the case in terms of the relationship between the number of connections of region has and the strength of those connections (middle). To ensure that we control for the analytical relationship between degree and strength, we also provide a version here the degree is controlled for in the strength measure (right). **b** In canonical idealized brain networks (left), regions with high levels of clustering have lower levels of betweenness centrality (yellow node) and vice-versa (light blue node). We show that in the best simulated networks, this is also the case in terms of the relationship clustering and betweenness (middle). We show this also for the weighted versions of the measure (right).

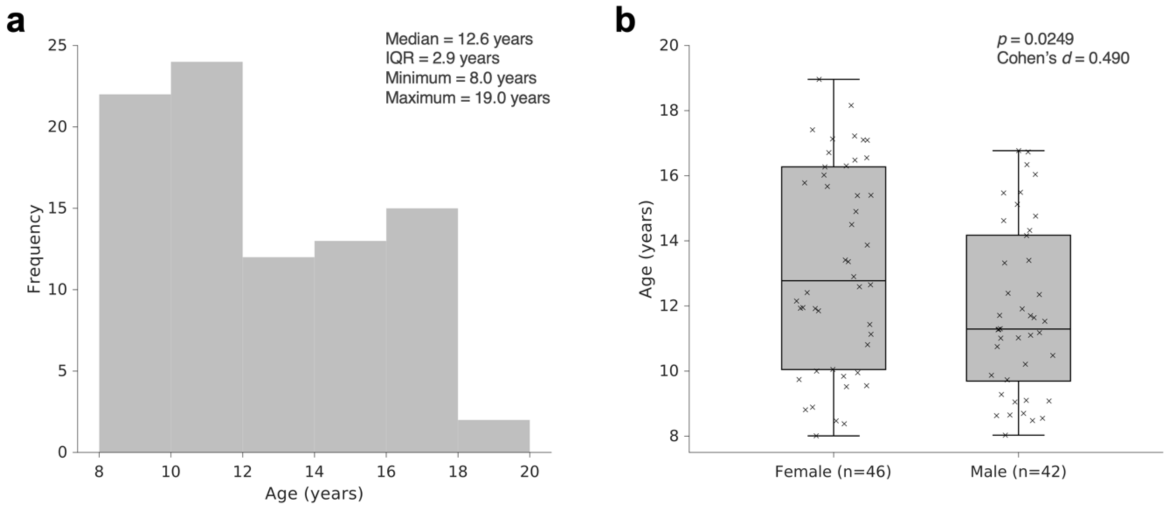


**Supplementary Fig. 4 | Sample age and sex characteristics. a** A histogram showing the age distribution across our $n$ = 88 sample. **b** A box plot showing age of females (left, $n$ = 46) and males (right, $n$ = 42). An interaction exists in our sample such that the females in our sample are slightly older than males.

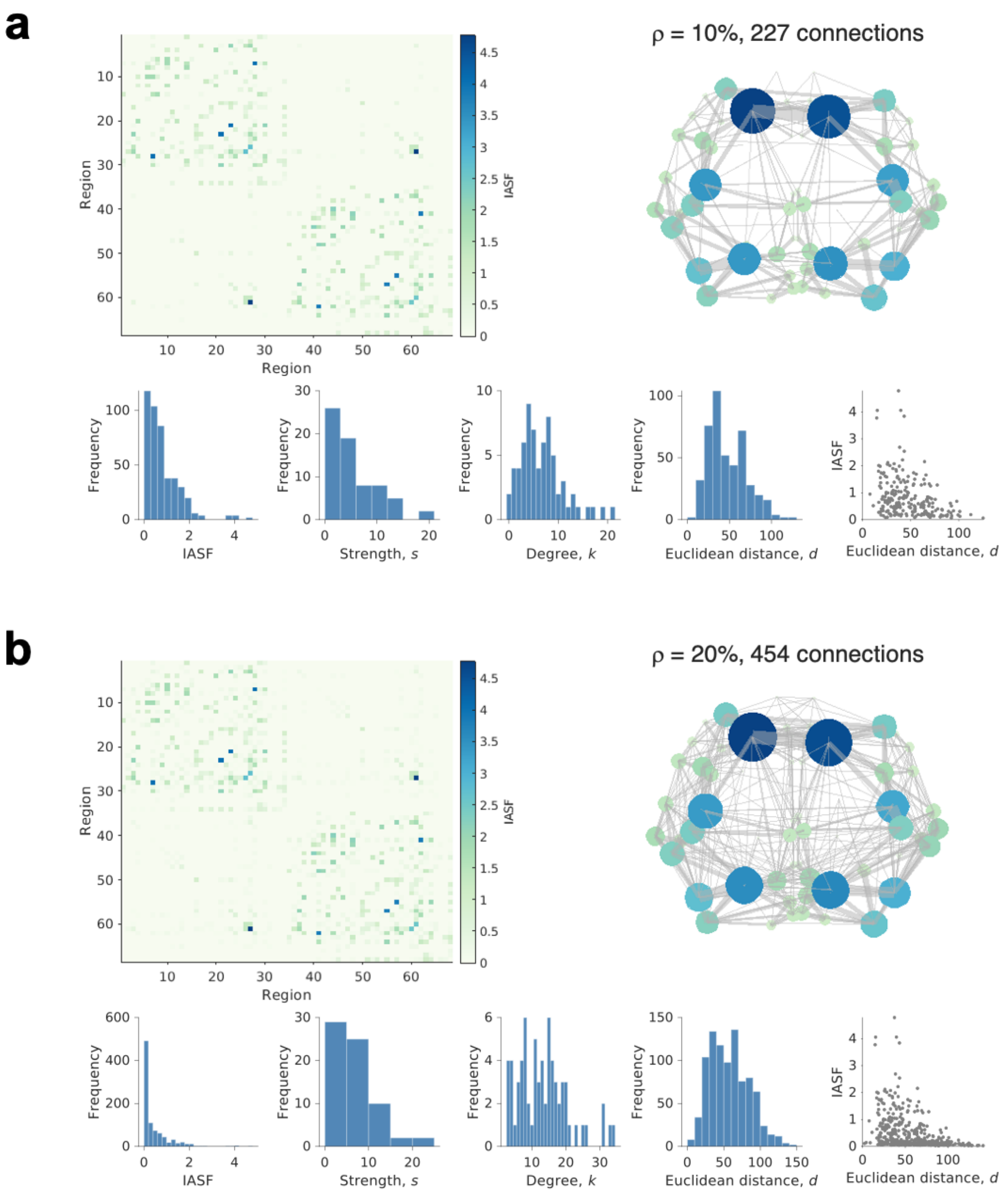


**Supplementary Fig. 5 | Empirical consensus network statistics. a** In sparse (ρ = 10% density networks, 227 connections) we show the empirical consensus weight matrix (top left), a graphical representation where edge width reflects intra-axonal signal fraction (IASF) and node size reflects node strength (top right). We then provide histograms of the empirical data across numerous measures (bottom). **b** We show the same in the dense (ρ = 20% density, 454 connections) networks.
